# DNA methylation cues in nucleosome geometry, stability, and unwrapping

**DOI:** 10.1101/2021.07.16.452533

**Authors:** Shuxiang Li, Yunhui Peng, David Landsman, Anna R. Panchenko

## Abstract

Cytosine methylation at the 5-carbon position is an essential DNA epigenetic mark in many eukaryotic organisms. Although countless structural and functional studies of cytosine methylation have been reported in both prokaryotes and eukaryotes, our understanding of how it influences the nucleosome assembly, structure, and dynamics remains obscure. Here we investigated the effects of cytosine methylation at CpG sites on nucleosome dynamics and stability. By applying long molecular dynamics simulations (five microsecond long trajectories, 60 microseconds in total), we generated extensive atomic level conformational full nucleosome ensembles. Our results revealed that methylation induces pronounced changes in geometry for both linker and nucleosomal DNA, leading to a more curved, under-twisted DNA, shifting the population equilibrium of sugar-phosphate backbone geometry. These conformational changes are associated with a considerable enhancement of interactions between methylated DNA and the histone octamer, doubling the number of contacts at some key arginines. H2A and H3 tails play important roles in these interactions, especially for DNA methylated nucleosomes. This, in turn, prevents a spontaneous DNA unwrapping of 3-4 helical turns for the methylated nucleosome with truncated histone tails, otherwise observed in the unmethylated system on several microsecond time scale.

## Introduction

DNA methylation plays an essential role in epigenetic signaling, specifically in regulating gene expression, genomic imprinting, X-chromosome inactivation, genome reprogramming (1), and abnormal methylation patterns are associated with several types of cancer and other diseases (2, 3). DNA methylation in human occurs primarily on the fifth carbon atom of cytosine within a CpG dinucleotide, forming 5-methylcytosine (5mC). It is estimated that about 80% of all CpG sites in the human genome are methylated (4). Methyl group on 5mC can directly or indirectly affect interactions with DNA-binding proteins, for example, it can hinder the binding of transcription factors (5) or mediate the binding of methyl-CpG-binding domains that recognize methylated CpG sites (6).

The effects of cytosine methylation on free DNA mechanical properties have been an issue of discussion for the past few years. Single-molecule atomic force microscopy experiments have shown that in the context of short DNA oligonucleotides, 5mC methylation results in either an increase or decrease in DNA structural stability in a manner that is sequence-dependent (7). A recent work using persistence length measurements demonstrates that cytosine methylation leads to longer contour lengths and increased DNA flexibility (8). Additionally, studies using infrared spectroscopy reveal that CpG methylation considerably shifts the equilibrium between different backbone sugar pucker conformations of DNA in solution (9). However, crystallographic experiments suggest that adding a methyl group to cytosine has negligible effects on the overall double-helical DNA structure (10).

There is plenty of evidence, although contradictory, that DNA methylation can influence the nucleosome assembly (11), its dynamics, stability (12–15), and positioning (16, 17). For example, recent single-molecule fluorescence studies relying on *in vitro* reconstituted nucleosomes and biochemical assays have revealed that DNA methylation promotes nucleosome compaction (12, 18) and increases the affinity of histones for DNA (11). However, subsequent studies present some evidence that cytosine methylation leads to a decreased level of compaction of nucleosomal DNA concluding that cytosine methylation probably causes nucleosome mechanical destabilization (13, 15). Moreover, X-ray and solid-state nanopore force spectroscopy at the level of mono-nucleosomes report that 5mC methylation does not perturb the structure of the nucleosome core particle and has no effect on nucleosome stability (19, 20). In addition to experimental studies, *in silico* approaches also yield contradictory results of the effects of cytosine methylation on DNA mechanics and nucleosome stability and dynamics. Although these simulations were performed on relatively short time scales of ~100 ns, they produced contrasting results that cytosine methylation stiffened DNA (21–23), increased the DNA molecule flexibility (24, 25), or had no effect (26).

Therefore, the underlying molecular mechanism of DNA methylation effects on the structure and stability of DNA, nucleosomes, and chromatin remains elusive and inconclusive. In this work, we investigated how 5mC methylation modulates the mechanical properties of nucleosomal and linker DNA and affects the nucleosome stability and dynamics. We succeeded in performing multiple all-atom molecular dynamics simulations on a relatively long five-microsecond time scale each, totaling in 60 microseconds. Our results showed that methylated cytosines at CpG sites significantly impacted the DNA geometry inducing undertwisting and underwinding of DNA, leading to a more extensive set of interactions between histones and DNA. This is turn, may explain the restricted capacity of the methylated DNA for spontaneous unwrapping which otherwise is observed for conventional nucleosomes without tails.

## Results

### DNA methylation prevents DNA unwrapping in nucleosomes

Using a high-resolution X-ray structure of the nucleosome core particle as the template and our previous protocol (27), we built an initial nucleosome model with the native DNA sequence of the *KRAS* gene that comprises 23 pairs of CpG sites (Figure S1) in total including 20 bp linker DNA flanking the core particle on each side (Figure 1A). The methylated nucleosome structure was obtained by converting all cytosines of CpG sites into 5mC (Figure 1B). Considering that the DNA dynamics can be restricted by histone tails, four structural models were constructed based on the combination of the CpG methylation status and presence or absence of histone tails: ^un^NUC_notail_, ^un^NUC_tail_, ^me^NUC_notail_, ^me^NUC_tail_ (Figure S2). All atom molecular dynamics (MD) simulations were then conducted for all models, and for each system three independent runs were performed, 12 runs in total, on a five-microsecond time scale each.

**Figure 1.**
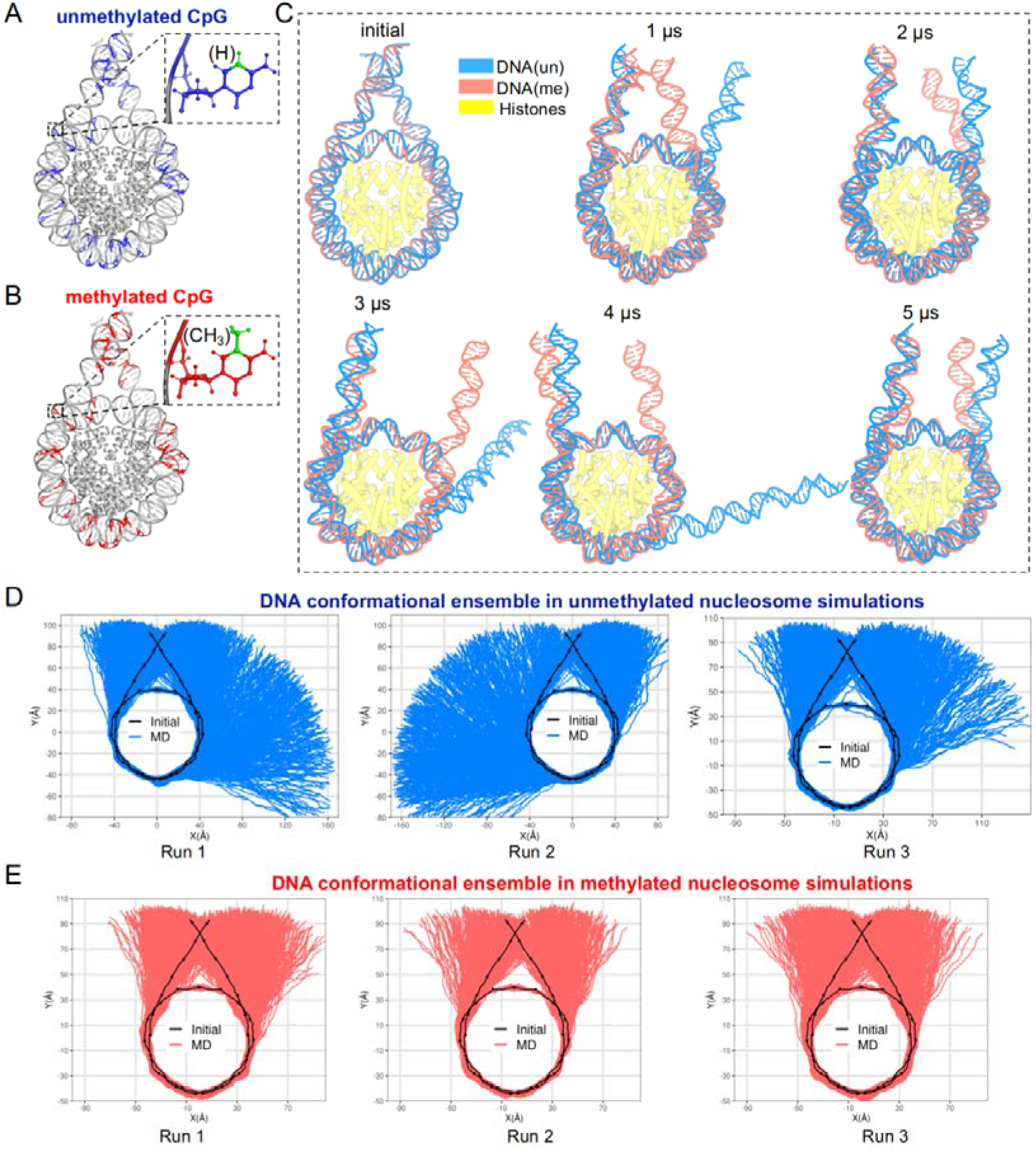
The dynamics of nucleosomal DNA in nucleosome systems without histone tails. (A, B) Illustrations of nucleosome systems with unmethylated (blue) and methylated (red) cytosines of the *KRAS* gene sequence (see Figure S1 for the full DNA sequence and methylated CpG sites in detail). (C) Nucleosome conformations of ^un^NUC_notail_ (blue) and ^me^NUC_notail_ (red) at 0 μs, 1 μs, 2 μs, 3 μs, 4 μs, and 5 μs simulation time points for two representative simulations. Two Run 1 simulations from Figure 1D and 1E are used for illustrating the representative conformations of ^un^NUC_notail_ (blue) and ^me^NUC_notail_ (red), respectively. (D) The DNA conformational ensembles in three independent ^un^NUC_notail_ simulation replicas, five microsecond each. (E) The DNA conformational ensembles in three independent ^me^NUC_notail_ simulation runs, five microsecond each. The black line and dots indicate the integer and half-integer SHL values of the initial conformation.

We first compare the MD trajectories of DNA in the ^un^NUC_notail_ and ^me^NUC_notail_ systems to examine the effect of CpG methylation on conformational changes of the nucleosomal DNA. The most conspicuous observation is the spontaneous unwrapping of several turns of the DNA ends from histone octamer in the ^un^NUC_notail_ simulations (Figure 1C). In all three simulation runs, the unwrapping process occurs asymmetrically at only one side of the nucleosome, at either entry or exit DNA sites (Figure 1D). As can be seen on Figure 2 and Figure S3, for the unmethylated nucleosome systems without tails, the pronounced DNA unwrapping starts after 2 to 3 μs extending up to SHL±4 with ~30 base pairs unwrapped from the octamer. There are several rewrapping events (1 to 3 occurrences) observed for all three simulation runs in the unmethylated systems without tails. This result is consistent with previous *in vitro* single-molecule FRET observations showing that the nucleosomal DNA unwraps asymmetrically under tension and unwrapping on one end stabilizes nucleosomes on another end (22). In contrast, the number of spontaneously unwrapped DNA base pairs in the methylated system is found to be less than ten, except for several frames exhibiting fast unwrapping and rewrapping motions encompassing of about 10-20 base pairs (Figure 2 and Figure S3). These results demonstrate that DNA methylation at CpG sites may affect the nucleosome dynamics and results in more stable and compact nucleosomes whose DNA ends are refractory to unwrapping. It is consistent with the previous FRET observations pointing to more compact methylated nucleosomes compared to the conventional unmethylated nucleosomes (12, 18). Such results are observed for all three ^me^NUC_notail_ simulation runs (Figure 1E).

**Figure 2.**
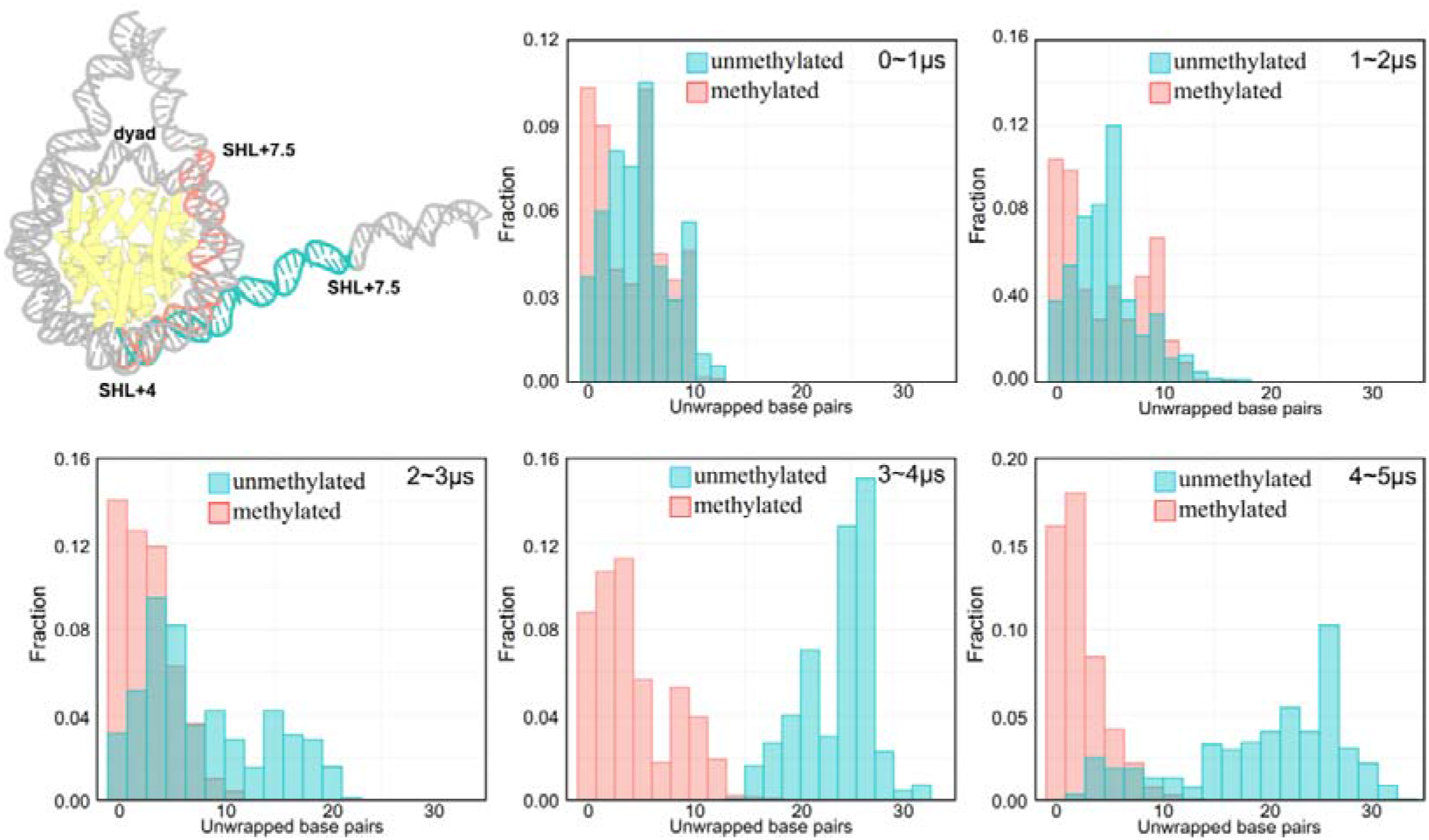
Time evolution of the number of unwrapped DNA base pairs during simulations for nucleosome systems without tails: ^un^NUC_notail_ (green) and ^me^NUC_notail_ (red). The number of unwrapped DNA base pairs were calculated based on the DNA sequence ranging from SHL+4 to SHL+7.5 (outer DNA region, see Figure S3A for definitions of DNA regions). The y-axis shows the fractions of frames with each individual unwrapped base pairs. Out of three ^un^NUC_notail_ simulation runs, one run with the most extent of DNA unwrapping from the exit side is shown. The simulation runs with the most extent of DNA unwrapping from the entry side is illustrated in Figure S3B.

It should be noted that all striking DNA unwrapping processes are observed only in the systems in which histone tails were truncated. Various studies previously demonstrated that histone tails considerably restricted the DNA breathing motions (28–30). Consistent with this, on 5 μs simulation timescale, both unmethylated and methylated nucleosomes with the full histone tails (^un^NUC_tail_ and ^me^NUC_tail_) do not yield substantial DNA unwrapping, only a transient detachment of a few base pairs from the histone octamer (Figure S4). However, as shown in the next section, both physical properties of DNA and histone-DNA interactions change markedly when methylated cytosines are incorporated for systems without and with histone tails.

### Cytosine methylation changes the DNA geometry

To investigate the molecular mechanism of how cytosine methylation may affect nucleosome stability, we start with the analysis of the local DNA geometrical and conformational properties for nucleosome systems with tails. We first characterize DNA conformations in terms of the base pair (shear, stretch, stagger, buckle, propeller and opening) and base-pair step (roll, tilt, twist, slide, shift and rise) parameters. Out of all DNA structural parameters, roll and twist values at the methylated sites show the most noticeable differences compared to CpG sites for unmethylated system. The distribution of twist values for the methylated steps is considerably shifted towards lower values (average value of twist is decreased by 7° per dinucleotide step, about 20% decrease) (Figure 3A). This result is corroborated by all six simulation runs. No significant change in twist parameters is observed for non-CpG sites between ^me^NUC_tail_ (red) and ^un^NUC_tail_ systems. To investigate the overall helical twist change of the double-strand DNA upon CpG methylation, we calculated the number of DNA turns from SHL-9 to SHL+9 which included the linker and nucleosomal DNA regions. The results show that methylated DNA is on average 0.4 turns underwound compared to unmethylated one (Figure 3E). We then determined the underwinding effect for the nucleosomal and linker DNA regions and found in both regions a reduced number of DNA turns in methylated nucleosomes (Figure S5). Therefore, we conclude that cytosine methylation results in undertwisting and underwinding of the DNA double helix, inducing negative stress on both the nucleosomal and linker DNA.

**Figure 3.**
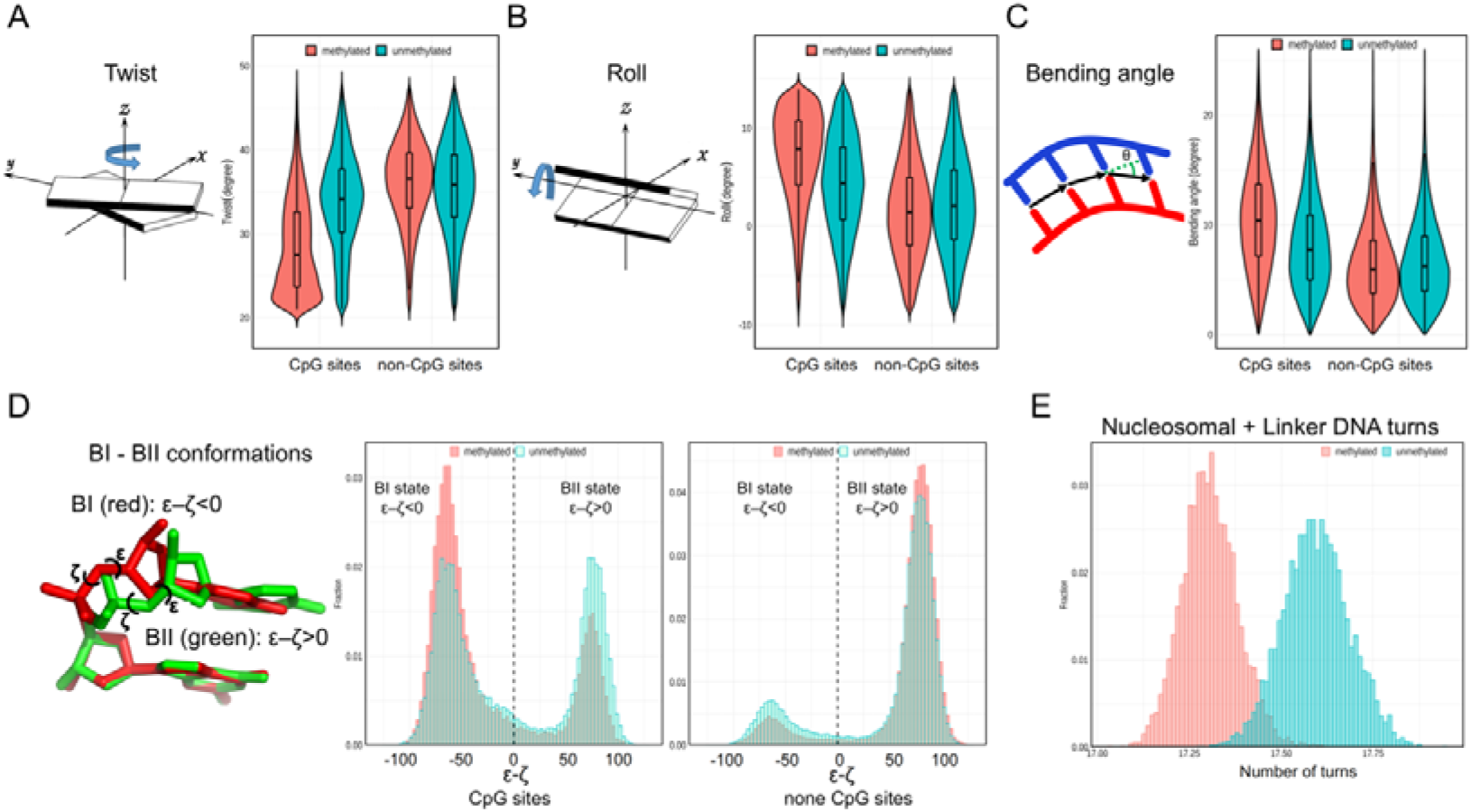
Effect of CpG methylation on DNA structural parameters. (A) Methylation reduces twist values at CpG sites. (B) Methylation increases roll values at CpG sites. (C) Methylation induces a more curved base-pair step with increased bending angles. (D) DNA sugar-phosphate backbone shifts the equilibrium toward the BI conformations at CpG sites upon methylation. (E) Methylation undertwists the overall DNA helical axis and induces decreased number of helical turns. The results for ^un^NUC_tail_ systems are shown in green and for ^me^NUC_tail_ (red) are shown in red.

Roll is another DNA structural parameter whose value changes considerably upon methylation (Figure 3B). At all CpG steps, roll values increase by an average of 3.5° upon methylation. This increment is consistent with the previous MD simulations of DNA oligomers that produced a similar effect size for roll parameters at methylated steps (31). Indeed, a systematic increase in bending angle values in all MD runs by 3° on average has been observed in methylated compared to unmethylated CpG steps (Figure 3C). It should be noted that no significant correlation is observed between different base-pair step parameters and bending angle values (Figure S6). Additionally, the overall curvature values for most regions of nucleosomal and linker DNA also increase in the methylated compared to the unmethylated systems (Figure S7). In previous experiments performed on DNA oligomers, it has been shown that DNA methylation can enhance DNA curvature (32). Our data are consistent with this observation, suggesting that the DNA methylation produces significant structural bending deformation. The effects of CpG methylation on all base pair and base-pair step parameters are summarized in Figure S8 and S9, respectively. It should be noted that the effects of cytosine methylation on DNA geometry are sequence dependent (Figure S10 for NUC_notail_ systems and Figure S11 for NUC_tail_ systems). The changes of these parameters also depend on whether the regions compose linker (free) DNA or nucleosomal DNA. The CpG steps of the linker DNA exhibit more prominent effects upon methylation than those from the nucleosomal DNA (Figure S12 and Figure S13).

Besides base-pair step parameters, it is important to characterize the conformation of the DNA sugar-phosphate backbone which is usually described by ε and ζ dihedral angles adopting either the canonical BI or BII configurations. As presented in Figure 3D, cytosine methylation shifts the equilibrium at CpG steps toward the BI conformational states and neighboring sites toward the BII conformations (see also Figure S14A and S14C). These BI/BII conformational change results in a shift toward the alternating BII-BI-BII conformation which has been previously shown to have the enhanced stability (33).

### Cytosine methylation enhances the DNA-histone interactions

Interactions between histones and DNA play essential roles in regulating the nucleosome dynamics and stability. We analyze these interactions and the first striking observation is that the overall number of contacts between histone and DNA is significantly increased upon DNA methylation for all histone types in different simulation runs (Figure 4A). The histone core-DNA interactions account for approximately 85% of the entire histone-DNA atom-atom contacts in both ^me^NUC_tail_ and ^un^NUC_tail_ systems (Figure 4B); the remaining contacts come from the interactions between DNA and histone tails (Figure 4C). A detailed inspection of the simulation trajectories reveals that the decreased conformational fluctuations of DNA in the methylated nucleosomes (Figure S17), especially DNA regions near the entry/exit sites, provide a potential pathway for DNA stabilization by histone core regions as well as by the H3 N-terminal and H2A C-terminal tails. We find from^me^NUC_tail_ simulations that the H2A C-terminal tail makes many stable contacts (see definition in Methods) with DNA near the entry/exit sites and forms partially compact secondary structure (Figure 4D, 4E). These DNA-histone tail interactions can stabilize the nucleosome and make the fluctuations at the very end of the nucleosomal DNA relatively small compared to the ^un^NUC_tail_ systems, which may suppress the DNA unwrapping upon methylation. Besides, since DNA-histone interactions mainly stem from the insertion of arginine residues into the DNA minor grooves (Figure S18), we examine these specific interactions in detail. We observe that the average number of contacts between the key arginines and DNA is dramatically increased in the methylated systems with tails (Figure 4G, Figure S19). These differences are especially pronounced for H2A R78 and H2B R36, which interact with the outer DNA turn, where the number of interactions almost doubled. Figure 4F shows the time evolution of the distances between H2A R78 and the DNA minor groove. The side chain from H2A R78 is consistently inserted into the DNA minor groove and rarely loses contacts in the methylated nucleosome systems. However, the contacts between H2A R78 and DNA are continuously disrupted in the unmethylated systems (Figure 4F, see Figure S20 for other simulations runs). Interestingly, the enhanced histone-DNA contacts in methylated nucleosomes are also observed for other residues, such as threonine and serine, but not for lysine residues (Figure S21). An increased number of contacts between methylated DNA and histones can explain the decreased DNA flexibility in the linker and nucleosome entry/exit sites (Figure S17). This observation is confirmed for both systems with and without histone tails. It contributes to the relatively stable (less prone to unwrapping) nucleosomes with methylated DNA compared to unmethylated ones, as shown in the previous section.

**Figure 4.**
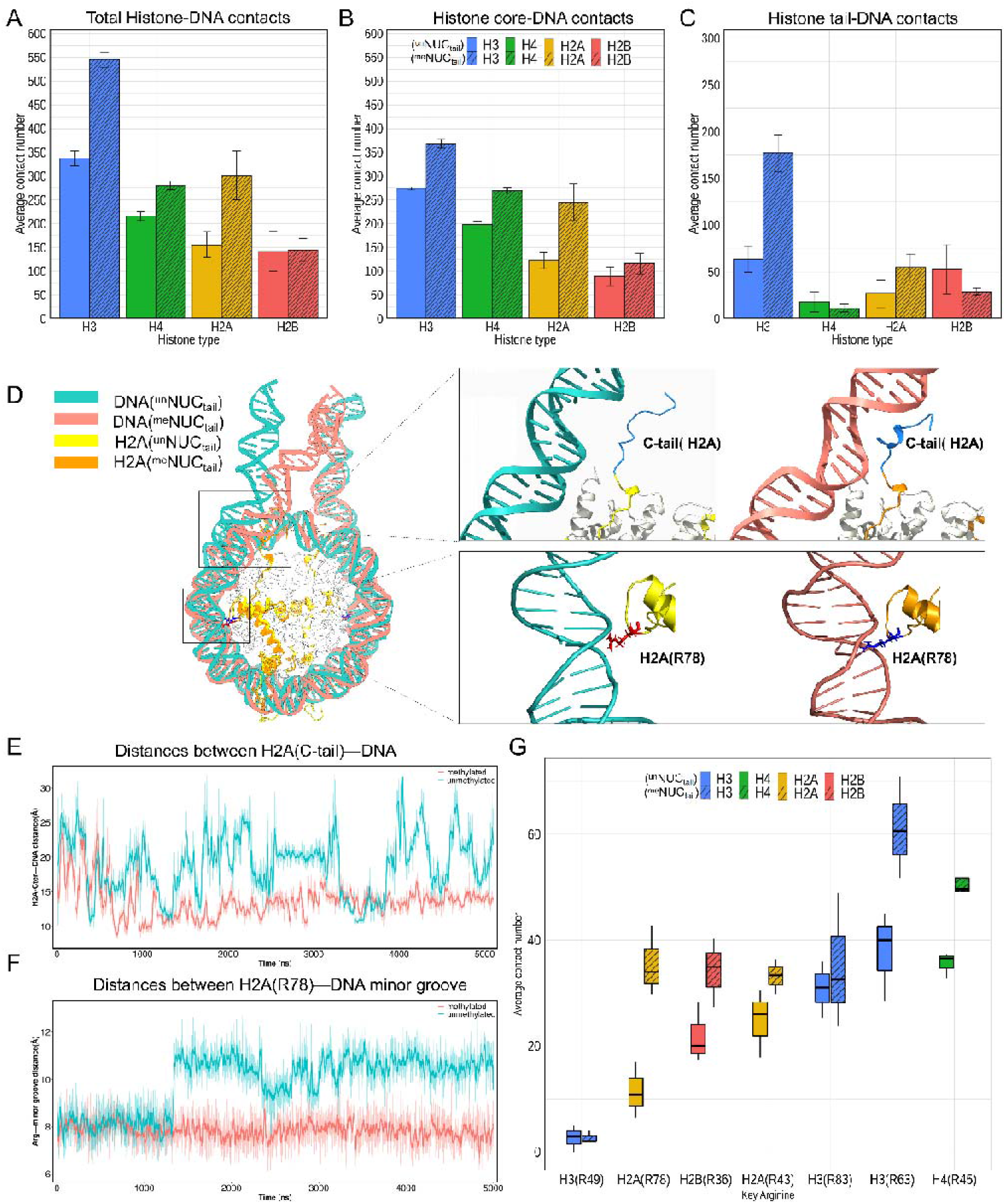
Atomic interactions between DNA and histones. (A-C) Average contact number of histone-DNA interactions for ^un^NUC_tail_ (without stripes) and ^me^NUC_tail_ (with stripes). (D) Representative snapshots showing the interactions between histones and DNA in ^me^NUC_tail_ (red) and ^un^NUC_tail_ (green) systems. The upper inset shows the interactions between H2A C-terminal tail with DNA and the lower inset shows interactions between H2A R78 and DNA. (E) Time evolution of the distances between the H2A C-terminal tail (center of residues 119 to 128) and the center of DNA segment (bp position from −65 to −73 relative to dyad) during the MD simulations. Thick lines were smoothed with Savitzky-Golay filter using ten ns window and first-degree polynomial. (F) Time evolution of the distances between the H2A(R78) and the DNA minor grooves (bp position from −58 to −54 relative to dyad) during the MD simulation. (G) Average contact number between DNA and the key arginines in the ^un^NUC_tail_ (without stripes) and ^me^NUC_tail_ (with stripes) systems. Error bars represent the standard errors calculated from two copies of each type of histone from DNA end three independent simulation runs.

## Discussion

Previous research presented insightful knowledge toward understanding of the impact of DNA methylation. Introducing hydrophobic 5mC groups in DNA which are pointed towards the major grooves may dramatically change geometry, dynamics and hydration patterns of DNA molecules (34). Nevertheless, the atomic mechanisms of the effects of DNA methylation on nucleosomes and chromatin structure and dynamics at the nucleosomal level remain ambiguous, if not contradictory (7, 31, 35). In addition, the DNA methylation effects *in vivo* vary considerably between organisms. In the present study, multiple runs of all-atom MD simulations have been performed to characterize the structural and dynamic differences between conventional and DNA methylated nucleosomes on several microsecond time scales. Our main results show that the changes in the geometrical properties of DNA and the interactions with histones affect nucleosome dynamics and stability. We show that 5mCpG leads to the enhanced bending at the CpG steps and is accompanied by changes in dinucleotide step parameters: the increased roll and decreased twist values compared to unmethylated systems. In regard to the sugar-phosphate backbone geometry, it was found that cytosine methylation shifts the population equilibrium of CpG steps toward the BI states due to methyl–sugar steric clashes (24). This, in turn, can enhance the intrinsic bending of DNA and reduce the stretching and twist stiffness as shown in both computational (24) and experimental studies (36).

Additionally, we observe the spontaneous unwrapping of unmethylated nucleosomal DNA exposing up to four superhelical turns for tailless nucleosomes in all simulation runs, but not for the nucleosome with the full histone tails. This result validates the importance of histone tails for maintaining the wrapped nucleosome state (30). It has been shown that the removal of histone tails can induce significant extent of DNA unwrapping from the histone octamer relative to the systems without histone tails (37–40). In our simulations, histone tails formed various interactions with the DNA ends to suppress DNA unwrapping in both the conventional and methylated nucleosomes. Moreover, the number of interactions formed by the H2A and H3 tails with DNA ends doubled and tripled respectively in the DNA methylated nucleosomes compared to the conventional unmethylated systems. This effect was not observed for the H4 and H2B histones. A recent work has demonstrated that destabilization of the H2A C-terminal tail in CENP-A nucleosomes facilitated DNA unwrapping (41). In addition, previous studies showed that the interactions between histone tails and the DNA ends arise primarily from the H3 and H2A tails. These interactions restrict the DNA end fluctuations and suppress further unwrapping of the nucleosomal DNA by inserting the H2A C-terminal tail into the minor groove and/or by accommodating the long H3 N-terminal tail between the DNA gyres (27, 38, 42). Moreover, enhanced key arginines-DNA interactions also promote the nucleosome stability and compactness in methylated systems compared to unmethylated ones. Namely, 10 out of 14 key arginines in the methylated nucleosomes are found to form more DNA-histone core contacts than those in the unmethylated systems. All these observations indicate an interesting possibility that the enhanced DNA interactions with the histone core regions as well as histone tails in DNA methylated nucleosomes, especially at the very end of the core DNA, suppress the DNA unwrapping from the histone octamer.

Based on our findings, the following mechanism emerges: changes in methylated DNA geometry might induce a more bent and under-twisted DNA to accommodate additional contacts between DNA and histone octamer, suppressing the DNA unwrapping. Our observations support the experimental data according to which positive torsional stress destabilizes nucleosomes and negative torsional stress, on the contrary, stabilizes nucleosome formation (43). In addition, a recent computational study has corroborated these findings by applying torsional stress to the nucleosomal DNA for tail-less nucleosomes and demonstrated that DNA unwraps more readily under positive torsional stress (44). It is further shown that *in vitro* negative supercoiling promotes nucleosome assembly, while positive supercoiling prevents it (45). According to our findings, CpG methylation suppresses the DNA unwrapping from histone core, which gives rise to a more compact nucleosome as compared with the conventional unmethylated nucleosome. This result is also in agreement with previous single molecule FRET experiments (12, 18), but not consistent with other experimental (13, 15) and computational observations (46). Additionally, our study reveals that overall histone-DNA interactions are greatly enhanced upon CpG methylation. Indeed, it has been previously reported that DNA methylation increases the DNA affinity for histone octamer (47), and enhances nucleosome occupancy (48).

In summary, we favor the interpretation that it is the combination of overall DNA physical properties upon methylation (twist, roll, bending angle, and BI-BII conformational equilibrium) that determines the conformational preference of DNA in such a way that it enhances the interactions with histone cores and tails to form more compact nucleosomes resistant to unwrapping. Although we used human DNA sequence in our study, the proposed effects might be extrapolated to other eukaryotic organisms. It should be mentioned that our analysis provides only one side of the story since, *in vivo,* the impact of DNA methylation is more diverse. In addition to directly influencing the DNA and nucleosome geometry, their mechanical properties and dynamics, the DNA methylation may recruit or occlude the binding of various specific proteins of the replication and transcription regulatory machineries.

## Materials and Methods

### Nucleosome modeling with the native DNA sequence and simulation protocol

The systems studied here consist of four full nucleosome structural models with each model comprising two straight 20 bp long DNA duplexes as linkers and the nucleosome core particle. The initial structural model of the nucleosome core particle was constructed using the high-resolution structure of the nucleosome core particle (PDB ID: 1KX5) (49) as a template and the 187 bp nucleosome sequence was taken from *Homo sapiens KRAS* proto-oncogene. This native sequence has 23 CpG sites and all of them were methylated on both DNA strands in the constructed methylated nucleosome system. The CpG sites are positioned at −91, −88, −83, −70, −63, −55, −44, −40, −38, −35, −20, −17, −14, 3, 16, 29, 34, 60, 67, 80, 83, 86, and 90 with respect to the dyad. The histone tails were clipped from the original 1KX5 structure and were linearly extended into the solvent symmetrically oriented with respect to the dyad axis. We used the initial nucleosome model to build unmethylated and methylated nucleosomes.

All MD simulations were performed with the package GROMACS version 2019.3 (50) using the Amber14SB force field for protein and OL15 for nucleic acid parameters (51, 52). 5-methylcytosines were modeled using parameters from Lankas et al. (53), which was originally derived for Amber parmbsc0 (54). Simulations were performed in explicit solvent using an Optimal Point Charge (OPC) water model, which was recently shown to reproduce water liquid bulk properties and to provide accuracy improvement in simulations of nucleic acids and intrinsically disordered proteins (55). In each simulation system, the initial structure model was solvated in a box with a minimum distance of 20 Å between the nucleosome atoms and the edges of the box. NaCl was added to the system up to a concentration of 150 mM. The solvated systems were first energy minimized using steepest descent minimization for 10,000 steps, gradually heated to 310 K over the course of 800 ps using restraints, and then equilibrated for a period of 1 ns. After that, the production simulations were carried out in the isobaric-isothermic (NPT) ensemble up to 5 μs, with the temperature maintained at 310 K using the modified Berendsen thermostat (velocity-rescaling) (56) and the pressure maintained at 1 atm using the Parrinello–Rahman barostat (57). A cutoff of 10 Å was applied to short-range non-bonded vdW interactions, and the Particle Mesh Ewald (PME) (58) method was used to calculate all long-range electrostatic interactions. Periodic boundary conditions were used. Covalent bonds involving hydrogens were constrained to their equilibrium lengths using the LINCS algorithm (59), allowing a 2.0 fs time step to be employed. Coordinates of the solutes were collected every 100 ps yielding a total of 50,000 frames for further analysis.

### Analysis of nucleosome dynamics

MD trajectory snapshots were first processed by performing a root mean square deviation (RMSD) fit of the C-α atoms of histone core (excluding the histone tail regions: residues 1–15 and 119–124 for H2A, 1–29 for H2B, 1–43 for H3, and 1–23 for H4) to the minimized structure of the nucleosome. The first 200 nanosecond frames of each simulation were treated as equilibration periods and were excluded from the analysis. The time courses of the base pair and base-pair step parameters were obtained by analyzing the nucleosomal DNA structure in each snapshot using the 3DNA software (60). The root mean square fluctuation (RMSF) values of nucleosomal DNA were calculated for backbone atoms at each time point of the trajectories using the Gromacs inbuilt tools g_rmsf. The local bending angle of a base pair step was calculated using 3D-DART (61). The first (1^st^) and last (187^th^) base pairs along the DNA sequence were not included in the analysis. In-house codes written in Python were developed to quantify the histone-DNA interactions defined by two non-hydrogen atoms from histone and DNA within a distance less than 4.5 Å. The atomic contacts between DNA and histone molecules that are present in more than 70% of trajectory frames are defined as stable contacts. We also compared the base pair and base-pair step parameters from current simulation trajectories with those from the simulations based on the same force field and water model but using a simulation protocol of AMBER 18 package (62). No apparent differences were found for these calculated parameters between the GROMACS and AMBER simulation protocols (Figure S15 and S16).

## Acknowledgements

SL and ARP were supported by the Department of Pathology and Molecular Medicine, Queen’s University, Canada. ARP is the recipient of a Senior Canada Research Chair in Computational Biology and Biophysics and a Senior Investigator Award from the Ontario Institute of Cancer Research, Canada. ARP and SL acknowledge the support of the Natural Sciences and Engineering Research Council of Canada (NSERC). YP and DL were supported by the Intramural Research Program of the National Library of Medicine at the U.S. National Institutes of Health. This study used the high-performance computational resources from the Compute Canada (https://docs.computecanada.ca) and the Biowulf clusters at the National Institutes of Health (https://hpc.nih.gov/systems).

## Notes

### Competing Interest Statement

The authors have declared no competing interest.

